# Dendritic localization of mRNA in *Drosophila* Mushroom Body Output Neurons

**DOI:** 10.1101/2020.09.03.281899

**Authors:** Jessica Mitchell, Carlas S. Smith, Josh Titlow, Nils Otto, Pieter van Velde, Martin Booth, Ilan Davis, Scott Waddell

## Abstract

Memory-relevant neuronal plasticity is believed to require local translation of new proteins at synapses. Understanding this process requires the visualization of the relevant mRNAs within these neuronal compartments. Here we used single-molecule fluorescence *in situ* hybridization (smFISH) to localize mRNAs at subcellular resolution in the adult *Drosophila* brain. mRNAs for subunits of nicotinic acetylcholine receptors and kinases could be detected within the dendrites of co-labelled Mushroom Body Output Neurons (MBONs) and their relative abundance showed cell-specificity. Moreover, aversive olfactory learning produced a transient increase in the level of *CaMKII* mRNA within the dendritic compartments of the γ5β′2a MBONs. Localization of specific mRNAs in MBONs before and after learning represents a critical step towards deciphering the role of dendritic translation in the neuronal plasticity underlying behavioural change in *Drosophila*.

## Introduction

Memories are believed to be encoded as changes in the efficacy of specific synaptic connections. Dendritic localization of mRNA facilitates specificity of synaptic plasticity by enabling postsynaptic synthesis of new proteins where and when they are required (Holt et al., 2019). Visualizing individual dendritically localized mRNAs in memory-relevant neurons is therefore crucial to understanding this process of neuronal plasticity.

Single molecule fluorescence *in situ* hybridization (smFISH) enables cellular mRNAs to be imaged at single-molecule resolution through the hybridization of a set of complementary oligonucleotide probes, each labelled with a fluorescent dye. Recent improvements in smFISH permit mRNA transcripts to be visualized in the dense heterogenous tissue of intact *Drosophila* brains (Long et al., 2017; Yang et al., 2017). Combining whole fly brain smFISH with neuron-specific co-labelling makes *Drosophila* an ideal model to investigate cell-specific mRNA localization and whether it is regulated in response to experience.

Olfactory learning in *Drosophila* depresses cholinergic synaptic connections between odor-specific mushroom body Kenyon Cells (KCs) and MBONs (Cohn et al., 2015; Handler et al., 2019; Hige et al., 2015; Owald et al., 2015; Perisse et al., 2016; Séjourné et al., 2011). This plasticity is driven by dopaminergic neurons whose presynaptic terminals innervate anatomically discrete compartments of the mushroom body, where they overlap with the dendrites of particular MBONs (Aso et al., 2010; Burke et al., 2012; Claridge-Chang et al., 2009; Lin et al., 2014; Liu et al., 2012). Dopamine driven plasticity is mediated by cAMP-dependent signalling and associated kinases such as Calcium/Calmodulin-dependent protein kinase II (CaMKII) and Protein Kinase A (PKA) (Boto et al., 2014; Handler et al., 2019; Hige et al., 2015; Kim et al., 2007; Qin et al., 2012; Tomchik & Davis, 2009; Yu et al., 2006; Zhang & Roman, 2013). Here we demonstrate localization of mRNAs in the 3D volumes of MBON dendrites by registering smFISH signals with co-labelled neurons using a custom image analysis pipeline. Moreover, we find that aversive learning transiently elevates dendritic *CaMKII* transcript levels within γ5β′2a MBONs.

## Results and Discussion

### mRNA localization in the intact adult *Drosophila* brain

Mammalian CaMKII mRNA is transported to neuronal dendrites, where it is locally translated in response to neuronal activity (Bagni et al., 2000; Miller et al., 2002; Ouyang et al., 1999). *Drosophila* CAMKII is critical for behavioural plasticity (Griffith, 1997; Malik et al., 2013) and is also thought to be locally translated (Ashraf et al., 2006). However, fly *CAMKII* mRNA has not been visualized in neurons. We therefore first hybridized *CaMKII* smFISH probes to whole mount brains and imaged the MB calyx (Figure 1A, 1B), a recognisable neuropil containing the densely packed dendrites of ~2,000 KCs and their presynaptic inputs from ~350 cholinergic olfactory projection neurons (Bates et al., 2020), using a standard spinning disk confocal microscope. To detect and quantify mRNA within the 3D volume of the brain, we developed a FIJI-compatible custom-built image analysis tool that segments smFISH image data and identifies spots within the 3D volume using a probability-based hypothesis test. This enabled detection of mRNAs with a false discovery rate of 0.05. CaMKII smFISH probes labelled 56 ± 5 discrete puncta within each calyx (Figure 1B, 1C). In comparison, smFISH probes directed to the α1 nicotinic acetylcholine receptor (nAChR) subunit labelled 33 ± 2 puncta in the calyx (Figure 1B, 1C). Puncta were diffraction limited and the signal intensity distribution was unimodal (Figure 1D-D′), indicating that they represent single mRNA molecules.

**Figure 1.**
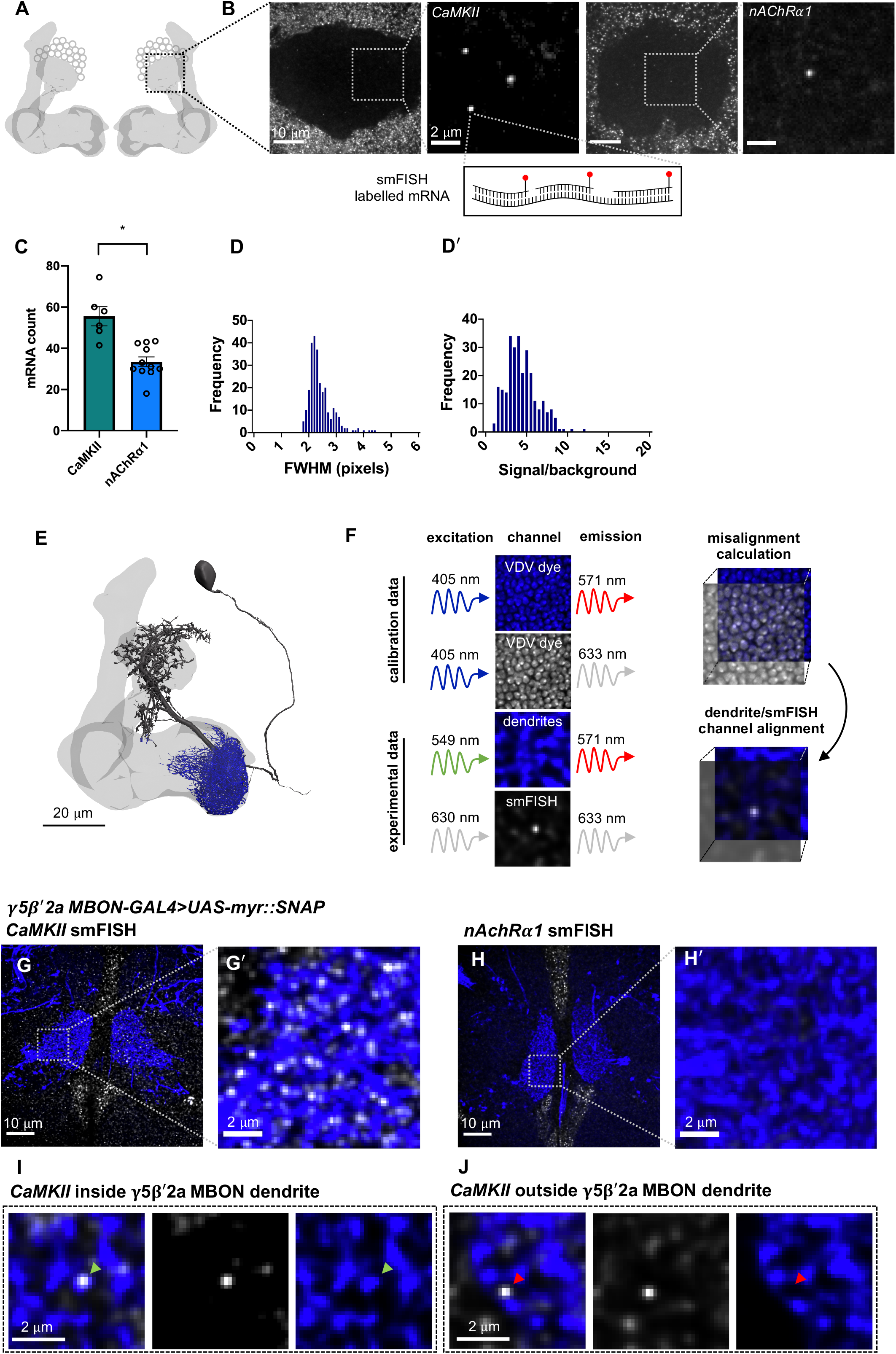
*CaMKII* and *nAChRα1* mRNA visualized in the MB calyx and γ5β′2a MBON dendrites with smFISH. **A.** Schematic of *Drosophila* MB. smFISH signal was imaged in the calyx, indicated by THE dashed box. **B.** *CaMKII* and *nAChRα1* mRNAs labeled with smFISH in the MB calyx. **C.** More *CaMKII* mRNAs are detected in the MB calyx relative to *nAChR 1 (*unpaired t-test: p=0.0003, t=4.727, df=15). **D.** smFISH spot size distribution (<u>F</u>ull <u>W</u>idth <u>H</u>alf <u>M</u>aximum, bottom) in MB calyx. **D′**. Unimodal smFISH spot intensity distribution (signal/background) indicates imaging at single-molecule resolution. **E.** Reconstruction of a γ5β′2a MBON (black) showing the dendritic field (blue) and MB (light grey). The projection to the contralateral MB is truncated. **F.** Alignment of dendrite and smFISH imaging channels using co-labeling with dsDNA VDV dye. VDV is excited with 405 nm and emission is collected in the dendritic and smFISH imaging channels, which were then aligned in x, y and z planes. **G-G′**. *CaMKII* smFISH within the γ5β′2a MBON dendrite co-labeled with R66C08-GAL4 driven UAS-UASmyr∷SNAP, and visualized with JF547SNAP dye. **H-H′**. nAchRα1 smFISH in γ5β′2a MBONs. **I.** Single *CaMKII* smFISH puncta localized within a γ5β′2a MBON dendrite (green arrowhead). **J.** Single *CaMKII* smFISH puncta localized outside of the γ5β′2a MBON dendrite (red arrowhead).

### mRNA localization within Mushroom Body Output Neuron Dendrites

*Drosophila* learning is considered to be implemented as plasticity of cholinergic KC-MBON synapses. To visualize and quantify mRNA specifically within the dendritic field of the γ5β′2a and γ1pedc>α/β MBONs we expressed a membrane-tethered UAS-myr∷SNAP reporter transgene using MBON-specific GAL4 drivers. This permitted simultaneous fluorescent labelling of mRNA with smFISH probes and the MBON using the SNAP Tag (Figure 1E). To correct for chromatic misalignment (Matsuda et al., 2018) that results from imaging heterogenous tissue at depth we also co-stained brains with the dsDNA-binding dye Vybrant DyeCycle Violet (VDV). VDV dye has a broad emission spectrum so labelled nuclei can be imaged in both the SNAP MBON and smFISH mRNA channels. This triple-labelling approach allowed quantification and correction of any spatial mismatch between MBON and smFISH channels in x,y and z planes, which ensures that smFISH puncta are accurately assigned within the 3D volume of the MBON dendritic field (Figure 1F).

Using this smFISH approach we detected an average of 32 ± 2 *CaMKII* mRNAs (Figure 1G-G′) within the dendrites of γ5β′2a MBONs. However, in contrast to the calyx, we did not detect *nAChRα1* in γ5β′2a MBON dendrites (Figure 1H-H′). This differential localization of the *CaMKII* and *nAChRα1* mRNAs within neurons of the mushroom body is indicative of cell-specificity. To probe mRNA localization in MBONs more broadly, we used a single YFP smFISH probe set and a collection of fly strains harboring YFP insertions in endogenous genes (Lowe et al., 2014). We selected YFP insertions in the *CaMKII*, *PKA-R2*, and *Ten-m* genes as test cases and compared the localization of their YFP-tagged mRNAs between γ5β′2a MBON and γ1pedc>α/β MBON dendrites.

The *CaMKII∷YFP* allele is heterozygous in flies also expressing myr∷SNAP in MBONs. Therefore, YFP smFISH probes detected half the number of *CaMKII* mRNAs in γ5β′2a MBON dendrites compared to *CaMKII*-specific probes (Figure 2A-A′, 2C). Importantly, YFP probes hybridized to YFP-negative control brains only produced a background signal (Figure 2B-B′) that was statistically distinguishable in brightness from genuine smFISH puncta (Figure 2D). These data indicate that the YFP probes have specificity and that the YFP insertion does not impede localization of *CaMKII* mRNA. We detected a similar abundance of *CaMKII∷YFP* in the dendritic field of γ1pedc>α/β MBONs. In contrast, we detected more *PKA-R2* mRNAs in γ5β′2a MBONs compared to γ1pedc>α/β MBONs. Surprisingly, *Ten-m* mRNAs were not detected in either γ5β′2a and γ1pedc>α/β MBON dendrites (Figure 2E, 2I), although they were visible in neighboring neuropil.

**Figure 2.**
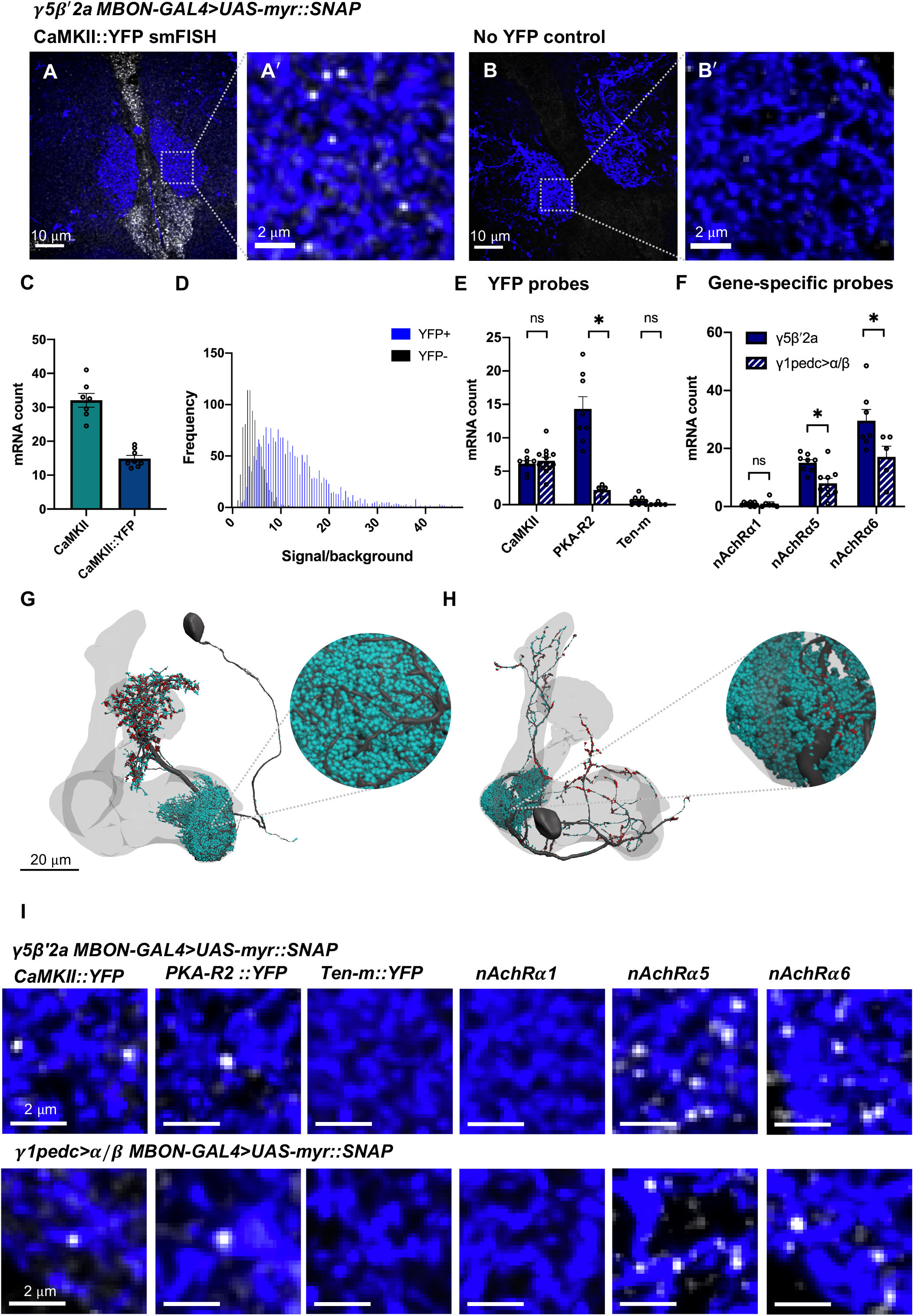
Differential localization of mRNAs in γ5β′2a and γ1pedc>α/β MBON dendrites. **A-A′.** *CaMKII∷YFP* mRNA visualized in γ5β′2a MBON dendrites using YFP smFISH probes. The γ5β′2a MBON is labelled by R66C08-GAL4 driven UAS-myr∷SNAP, and visualized with JF547SNAP dye. **B-B′**. YFP smFISH signal in a γ5β′2a MBON in a negative control fly. **C.** The *CaMKII∷YFP* allele is heterozygous resulting in detection of half as many *CaMKII* mRNAs in γ5β′2a MBONs using YFP probes relative to that detected with *CaMKII* gene-specific probes. **D.** smFISH puncta detected using YFP probes in *CaMKII∷YFP* brains have statistically distinguishable signal/background intensity distribution relative to puncta in control genotypes (Wilcoxon rank sum p<0.005). **E.** Quantification of mRNA localization in transcripts localize within the dendrites of γ5β′2a MBONs relative to γ1pedc>α/β MBONs (unpaired t-test: p=0.004, t=5.069, df=11). *Ten-m* mRNAs did not localize to either MBON dendritic field. *CaMKII* mRNAs were detected in equal abundance. **F.** Quantification of mRNA localization in γ5β′2a and γ1pedc>α/β MBON dendrites with gene-specific smFISH probes. *nAchRα1* mRNAs did not localize to the dendrites of either γ5β′2a or γ1pedc>α/β (unpaired t-test: p=0.046, t=2.274, df=10) mRNAs localized to γ5β′2a MBON dendrites relative to γ1pedc>α/β MBON dendrites. **G.** Reconstruction of a γ5β′2a MBON. Individual postsynapses (turquoise spheres) and presynapses (red spheres) are labelled. The projection to the contralateral MB is truncated. **H.** Reconstruction of a γ1pedc>α/β MBON. Individual postsynapses (turquoise spheres) and presynapses (red spheres) are labelled. The projection to the contralateral MB is truncated. **I.** Example smFISH images of mRNAs localized in γ5β′2a (R66C08-GAL4>UAS-myr∷SNAP) and γ1pedc>α/β MBON (MB112C-GAL4>UAS-myr∷SNAP) dendrites. Asterisks denote significant difference (p<0.05). Data are means standard error of mean (S.E.M.). Individual data points are displayed.

Although we did not detect *nAChRα1* mRNA within γ5β′2a MBON dendrites, prior work has shown that nAChR subunits, including nAChRα1, are required in γ5β′2a MBON postsynapses to register odor-evoked responses and direct odor-driven behaviours (Barnstedt et al., 2016). Since the YFP insertion collection does not include nAChR subunits, we designed *nAChRα5* and *nAChRα6* specific smFISH probes. These probes detected *nAChRα5* and *nAChRα6* mRNAs within γ5β′2a and γ1pedc>α/β MBON dendrites, with *nAChRα6* being most abundant (Figure 2F, 2I). The selective localization of *nAChRα5* and *nAChRα6* mRNA to MBON dendrites indicates that these receptor subunits may be locally translated to modify the subunit composition of postsynaptic nAChR receptors.

Localized mRNAs were on average 2.8x more abundant in γ5β′2a relative to γ1pedc>α/β MBON dendrites (Figure 2E, 2F). We therefore tested whether this apparent differential localization correlated with dendritic volume and/or the number of postsynapses between these MBONs. Using the recently published electron microscope volume of the *Drosophila* ‘hemibrain’ (Xu et al., 2020) (Figure 2G, 2H), we calculated the dendritic volume of the γ5β′2a MBON to be 1515.36 nm^3^ and the γ1pedc>α/β MBON to be 614.20 nm^3^. In addition, the γ5 and β′2a regions of the γ5β′2a MBON dendrite contains 30,625 postsynapses whereas there are only 17,020 postsynapses in the γ1 region of the γ1pedc>α/β MBON. Larger dendritic field volume and synapse number is therefore correlated with an increased number of localized mRNAs, suggesting that these parameters may be important determinants of localized mRNA copy number.

### Learning transiently changes *CAMKII* mRNA abundance in γ5β′2a MBON dendrites

We tested whether *CaMKII∷YFP* mRNA abundance in γ5β′2a and γ1pedc>α/β MBONs was altered following aversive learning (Figure 3A, 3B). We also quantified mRNA in the soma and nuclei of these MBONs (Figure 3A′, 3B′). Transcriptional activity is indicated by a bright nuclear transcription focus (Figure 3C). We subjected flies to four conditions (Figure 3D): 1) an ‘untrained’ group that were loaded and removed from the T-maze but not exposed to odors or shock. 2) an ‘odor only’ group, exposed to the two odors as in training but without shock. 3) a ‘shock only’ group that were handled as in training and received the shock delivery but no odor exposure. 4) a ‘trained’ group that were aversively conditioned by pairing one of the two odors with shock. Fly brains were extracted 10 min, 1 h or 2 h after training and processed for smFISH.

**Figure 3.**
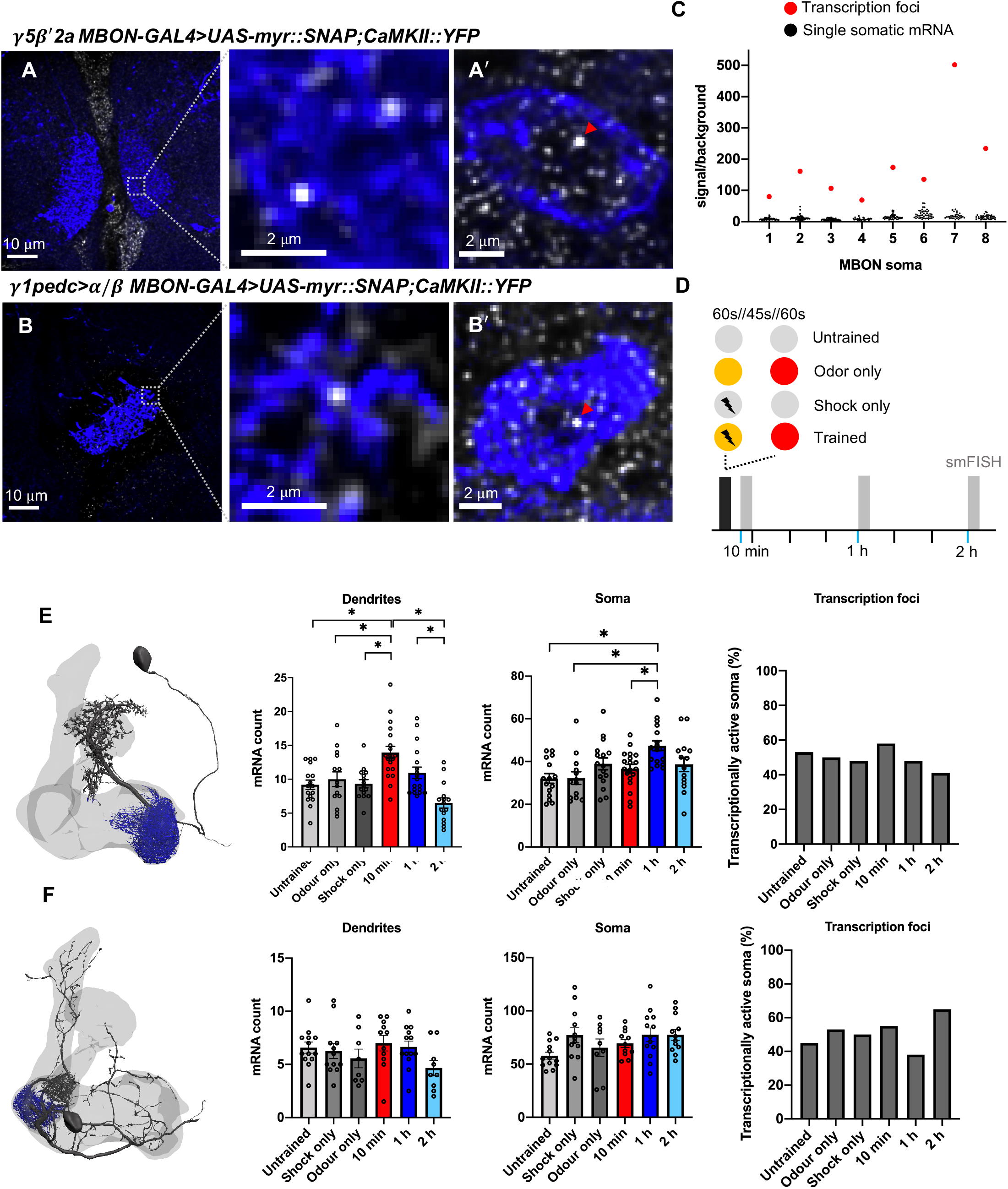
Learning alters CaMKII mRNA abundance in the γ5β′2a MBONs. **A-A′**. *CaMKII∷YFP* smFISH in γ5β′2a MBON dendrites and soma (R66C08-GAL4>UAS-myr∷SNAP). **B-B′**. CaMKII∷YFP smFISH in γ1pedc>α/β MBON dendrites and soma (MB112C-GAL4>UAS-myr∷SNAP). Nuclear transcription foci are indicated (red arrowheads). **C.** *CaMKII∷YFP* smFISH signal/background in γ5β′2a soma. Transcription foci are readily distinguished as the brightest puncta in the soma/nucleus (red data points). Note, only one transcription focus can be visualized per cell since the *CaMKII∷YFP* allele is heterozygous. **D.** Schematic of aversive training and control protocols followed by smFISH. **E.** *CaMKII∷YFP* mRNA numbers in γ5β′2a MBON dendrites increase 10 min after odor-shock pairing, relative to control groups (one-way ANOVA: untrained-10min p=0.001; odor only-10min p=0.016; shock only-10min p=0.002), and decrease to baseline by 2h (one-way ANOVA: 10min-2h p<0.001; 1h-2h p=0.004). *CaMKII∷YFP* mRNA numbers in γ5β′2a MBON soma increase 1 h after odor-shock pairing, relative to untrained (one-way ANOVA: p=0.001), odor only (one-way ANOVA: p=0.002), and 10 min post training (one-way unchanged (*X*^2^=2.064, df=5, p=0.840). **F.** *CaMKII∷YFP* mRNA numbers are not changed by p=0.212), their soma (one-way ANOVA: f=2.183, p=0.067), and there is no detected change difference (p<0.05). Data are means ± standard error of mean (S.E.M.). Individual data points are displayed.

*CaMKII* mRNA increased significantly in γ5β′2a MBON dendrites 10 min after training (Figure 3E) compared to all control groups. Levels returned to baseline by 1 h and remained at that level 2 h after training (Figure 3E). *CaMKII* mRNAs in γ5β′2a MBON soma showed a different temporal dynamic, with transcripts peaking 1 h after training, albeit only relative to untrained and odor only controls (Figure 3E). The proportion of γ5β′2a nuclei containing a *CaMKII* transcription focus did not differ between treatments (Figure 3E), suggesting that the transcript increase in the soma is not correlated with the number of actively transcribing γ5β′2a nuclei, at least at the timepoints measured. Assessing *CaMKII* mRNA abundance in γ1pedc>α/β MBONs after learning did not reveal a change in mRNA abundance in the dendrites or soma between trained flies and all control groups at all timepoints measured (Figure 3F). These results indicate specificity to the response observed in the γ5β′2a MBONs.

Since CaMKII protein is also labelled with YFP in *CaMKII∷YFP* flies, we also assessed protein expression by measuring YFP fluorescence intensity specifically within the MBON dendrites. This analysis did not reveal a significant difference in fluorescence intensity across treatments (Supplementary Figure 1). Since smFISH provides single-molecule estimates of mRNA abundance, a similar level of single-molecule sensitivity may be required to detect subcellular resolution changes in protein copy number.

Prior studies have demonstrated a learning-related increase in the response of γ5β′2a MBONs to the shock-paired odor and suggested this arises from release of feedforward inhibition from γ1pedc>α/β MBONs (Bouzaiane et al., 2015; Owald et al., 2015). We speculate that the change in *CaMKII* mRNA abundance might be a consequence of network-level potentiation of the activity of the γ5β′2a MBON, such as that that would result from a release from inhibition.

## Supporting information

Key Resources Table

## Acknowledgements

We are grateful for the microscopy facilities and expertise provided by Micron Advanced Bioimaging Unit (supported by Wellcome Strategic Awards 091911 and 107457). We thank Jeff Lee for assistance with smFISH probe generation and members of the Waddell group for discussion. J.M. was funded through the BBSRC Interdisciplinary Bioscience Doctoral Training Programme. C.S.S. and P.V.V. were funded by the Netherlands Organisation for Scientific Research (NWO), under NWO START-UP project no. 740.018.015 and NWO Veni project no. 16761. C.S.S. acknowledges a research fellowship through Merton College, Oxford, UK. I.D. was supported by a Wellcome Trust Senior Research Fellowship (096144), Wellcome Trust Investigator Award (209412), and Wellcome Trust Strategic Awards (091911 and 107457). S.W. was funded by a Wellcome Principal Research Fellowship (200846/Z/16/Z), an ERC Advanced Grant (789274), and a Wellcome Collaborative Award (203261/Z/16/Z).

## Author Contributions

Conceptualization: J.M., C.S.S., S.W. Investigation: J.M., C.S.S., N.O., P.V.V., J.T. Methodology: J.M., C.S.S., J.T. Resources: S.W., I.D. Software: C.S.S., P.V.V. Funding acquisition: S.W., I.D., M.B. Supervision: S.W. Writing – original draft: J.M., S.W. Writing – review & editing: S.W., J.M., C.S.S., N.O., J.T., I.D.

---

**Supplementary Figure 1. CaMKII∷YFP fluorescence intensity inγ5β′2a and γ1pedc>α/β MBONs after learning.**

**A.** CaMKII∷YFP fluorescence intensity (adu/voxel) within γ5β′2a MBON dendrites and soma.

**B.** CaMKII∷YFP fluorescence intensity (adu/voxel) within γ1pedc>α/β MBON dendrites and soma. No significant differences between trained and control groups were observed (One-way ANOVA/Kruskal Wallis p>0.05).

## Materials and Methods

### Fly Strains

Flies were raised on standard cornmeal agar food in plastic vials at 25 °C and 40-50 % relative humidity on a 12 h: 12 h light: dark cycle. Details of fly strains are listed in the Key Recourses Table.

### smFISH probes

Oligonucleotide probe sets were designed using the web-based probe design software https://www.biosearchtech.com/stellaris-designer. The YFP smFISH probe set was purchased from LGC BioSearch Technologies (California, USA) prelabelled with Quasar-670 dye. *CaMKII*, *nAChRα1*, *nAChRα5* and *nAChRα6* DNA oligonucleotide sets were synthesised by Sigma-Aldrich (Merck) and enzymatically labelled with ATTO-633 according to (Gaspar et al., 2017). DNA oligonucleotide sequences for each smFISH probe set are provided in the Supplementary Information.

### Whole *Drosophila* brain smFISH

Whole adult brain smFISH was performed essentially as described (Yang et al., 2017). 2-4 day old adult *Drosophila* brains were dissected in 1X phosphate buffered saline (PBS) and fixed in 4% v/v paraformaldehyde for 20 min at room temperature. Brains were washed 2X with PBS, followed by 20 min in 0.3% v/v Triton X-100 in PBS (PBTX) to permeabilise the tissue, then 15 min in PBTX with 500 nM JF549-SNAPTag (Grimm et al., 2015) for neuronal labelling. 3 × 10 min washes in PBTX removed excess dye. Samples were then incubated in wash buffer (2x RNase-free SSC + 10% v/v deionised formamide) for 10 min at 37 °C, wash buffer was replaced with hybridization buffer (2x RNase-free SSC, 10% v/v deionised formamide, 5% w/v dextran sulphate, 250nM smFISH probes) and samples incubated overnight at 37 °C. Hybridization buffer was removed before samples were washed 2X in freshly prepared wash buffer and incubated 40 min in wash buffer containing Vybrant DyeCycle Violet Stain (1:1000) to label nuclei. Samples were then washed 3X times in wash buffer, mounted on a glass slide covered with Vectashield anti-fade mounting medium (refractive index 1.45) and immediately imaged.

### Olfactory conditioning

Aversive olfactory conditioning was performed essentially as described by (Tully & Quinn, 1985). 3-octanol (OCT) was used as the shock-paired odor. 4-methylcyclohexanol (MCH) was used as the unpaired odor. Odors were prepared at concentrations of 9 μl OCT in 8 ml mineral oil, and 8 μl MCH in 8 ml mineral oil. Groups of ~100 flies were aliquoted into plastic vials containing standard cornmeal agar food and a 2 × 6 cm piece of filter paper. Flies were conditioned as follows: 1 min OCT paired with 12 × 90 V shocks at 5 s interstimulus interval; 45 s clean air; 1 min MCH. Control groups were handled in the same way except for the differing presentation of either odors or shock. Untrained flies experienced no odor or shock, the odor only group experienced the two odor presentations without shock, and the shock only group received the shock presentations but no odors. Aversive olfactory conditioning was performed at 23 °C and 70% relative humidity. Following training, flies were returned to food vials and brains were dissected either 10 min, 1 h or 2 h later, and smFISH analyses performed.

### Microscopy

Samples were imaged on a Spinning Disk confocal microscope (Perkin Elmer UltraView VoX) with a 60x 1.35 N.A. oil immersion UPlanSApo objective (Olympus) and a filter set to image fluorophores in DAPI, FITC, TRITC, and CY5 channels (center/bandwidth; excitation: 390/18, 488/24, 542/27, 632/22 nm; emission: 435/48, 594/45, 676/34 nm), the corresponding laser lines (488/4.26, 561/6.60, 640/3.2, 405/1.05, 440/2.5, 514/0.8 nm/mW) and an EMCCD camera (ImagEM, Hamamatsu Photonics). The camera pixel size is 8.34 μm, resulting in a pixel size in image space of approximately 139 nm. Optical sections were acquired with 200 nm spacing along the z-axis within a 512 × 512 pixel (71.2 × 71.2 μm) field of view.

### Deconvolution

Deconvolution was carried out using commercially available software (Huygens Professional v19.10.0p1, SVI Delft, The Netherlands). Raw image data generated in .mvd2 file format were converted to OME.tiff format using FIJI (Schindelin et al., 2012) (convert_mvd2_to_tif.ijm). Spherical aberration was estimated from the microscope parameters (see Microscopy). Z-dependent momentum preserving deconvolution (CLME algorithm, theoretical high-NA PSF, iteration optimized with quality change threshold 0.1% and iterations 40 maximum, signal to noise ratio 20, area radius of background estimation is 700 nm, a brick mode is 1 PSF per brick, single array detector with reduction mode SuperXY) was then applied to compensate for the depth-dependent distortion in point spread function thereby reducing artefacts and increasing image sharpness.

### Multi-channel alignment

Misalignment between channels was corrected for using Chromagnon (v 0.81) (Matsuda et al., 2018). To estimate channel misalignment nuclei were labelled with the broad emission spectrum dye (Vybrant DyeCycle Violet Stain, ThermoFisher) (Smith et al., 2015). The dye was excited at 405nm and emission was recorded using the appropriate filters for each imaging channel. Chromatic shift was estimated by finding the affine transformation that delivers a minimum mean square error between the nuclear stain in the various channels. Nuclear calibration channels for chromatic shift correction were separated using ImageJ (see macro Split_ometiff_channels_for_chromcorrect.ijm). The affine transformation was estimated and alignment was performed by calling Chromagnon from Python (see script chromagnon_bash.py). The resulting aligned and deconvolved images were saved in .dv format for further downstream analysis.

### Calculating postsynaptic abundance and volume of γ5β′2a and γ1pedc>α/β MBON dendrites

Neuro-morphological calculations were performed with navis 0.2.0 library functions in Python (https://pypi.org/project/navis/) (Bates et al., 2020) using data obtained from the *Drosophila* hemibrain dataset (v1.1) (https://neuprint.janelia.org) (Xu et al., 2020). To calculate the dendritic volume and postsynaptic abundance of γ5β′2a and γ1pedc>α/β MBONs, neuron skeletons, neuropil meshes, and synapse data were first imported. Neural skeletons were then used to generate 3D neuron reconstructions. Dendritic processes of the γ5β′2a MBON were determined by intersecting neuronal skeletons with the MB mesh containing the γ5 and β′2a compartments. Dendritic processes of the γ1pedc>α/β MBON were determined by intersecting the skeleton within the γ1 MB compartment mesh. The available γ1 MB compartment mesh did not encompass the entirety of the γ1pedc>α/βMBON dendrites in the γ1 MB compartment so the volume of the mesh was scaled up 1.35x. This intersects with almost all γ1pedc>α/β MBON dendrites in the γ1 MB compartment, but not any other substantial part of the neuron. Dendritic volume (nm^3^) was calculated as the sum of the neurite voxels multiplied by 8^3^ since the resolution of each voxel is 8 cubic nanometers. The number of postsynapses within these compartments was also determined using the synapse data that accompany the neuron skeletons (Xu et al., 2020).

### Data visualization

smFISH data were visualized in FIJI (Schindelin et al., 2012). Maximum intensity projections representing 2 μm sections are presented for visualization purposes. Figures 2E and 2F are single z-sections (representing a 0.2 μm section). 3D reconstructions of γ5β′2a and γ1pedc>α/β MBONs were created in Blender v.2.8.2 with navis 0.2.0 plugin and using data obtained from www.neuprint.janelia.org (Xu et al., 2020).

### mRNA detection

An smFISH spot detection Matlab script based on (Smith et al., 2015) was written to quantify localized mRNA transcripts in *Drosophila* brains. Software for processing smFISH datasets is available as Supplementary Software. Updates will be made at https://github.com/qnano/smfish. The smFISH channel was extracted and stored as a 3D greyscale image. mRNA signal was detected using 3D generalized likelihood ratio test (Smith et al., 2015).The false detection rate is 0.05 and the spot width is σ_x,y_ = 1.39 and σ_z_ = 3.48. After 3D detection, the intensity, background, width and subpixel position of the detected mRNA spots are estimated using maximum likelihood estimation (MLE) (Smith et al., 2010).To reduce the impact of overlapping spots in 3D, only a 2D cross-section is used from the z-plane where the spot is detected. To filter out spurious detections all spots with a width greater than 5 pixels are discarded.

### mRNA-dendrite co-localization

To quantify calyx and dendritic localized smFISH puncta, the calyx and dendritic area were first segmented manually. The contour of the calyx and dendritic area is converted to a mask (M_1_) using the MATLAB R2019b function roipoly. To quantify smFISH puncta co-localizing with dendrite label, a mask of the dendrite label is created by enhancing the image using a difference of Gaussians filter (width of 1 and 5 pixels) and then thresholding the product between the enhanced image (*A*) and masked area (M_1_) to obtain a mask (M_2_):

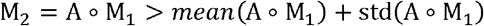

where mean() and std() are the sample mean and sample standard deviation of the image intensity values and *A ∘ B* is the Hadamard product between A and B. The sample standard deviation is calculated as:

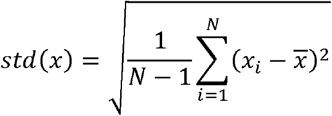

where *N* is the number of data points. smFISH signal within γ5β′2a MBON dendrites innervating the γ5 and β′2a MB compartments were analyzed. smFISH signal within the γ1pedc>α/β dendrites innervating the γ1 MB compartment were analyzed. Sections of 10 × 0.2 μm individual z-slices of MB calyx, γ5β′2a MBON dendrites, or γ1pedc>α/β MBON dendrites were analyzed. smFISH puncta overlapping with the calyx or dendrite mask were considered co-localizing and therefore localized within that neuronal compartment.

### Spot brightness and full width half maximum (FWHM) analysis

For each detection, a region of interest (ROI) is extracted as a 2D box in the x-y plane with a size of *2×(3*σ*_x,y_+1).* For each ROI the MLE of the × and y position, the number of photons, the number of background photons, and the width of the 2D Gaussian, σ*x,_y_* is computed. The FWHM of the spots is calculated as 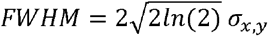

### Verification of transcription foci

Soma containing bright nuclear transcription foci were selected to quantify the difference in intensity relative to diffraction-limited smFISH puncta. The nuclear localization of the smFISH puncta with the highest photon count was validated by visual inspection and was considered to correspond to the transcription site. The width (σ_*x,y*_) of the transcription foci significantly differ from the sparse smFISH signal and is estimated by fitting a 2D Gaussian to the transcription site using the MATLAB 2019b non-linear least-squares routine *lsqcurvefit*. Transcription foci brightness and background were computed using the same MLE protocol as for diffraction-limited spots, but with the estimated σ_*x,y*_.

### YFP fluorescence intensity

To quantify YFP fluorescence intensity within co-labelled neurons, we developed a FIJI compatible macro plugin. Depth-dependent bleaching was first corrected for over the z-stack using an exponential fit. Background signal was then subtracted in each z-section using a rolling ball filter with a width of 60 pixels. 5 z-sections above and below the centre of the image were cropped for analysis. YFP fluorescence intensity was recorded within the dendrites or soma of the co-labelled neuron using the mask described above (mRNA-dendrite co-localization). Fluorescence intensity was calculated as analog digital units (adu)/volume (dendrites or soma) to give adu/voxel. Software for analysing fluorescent protein expression in single neurons is available as Supplementary Software. Updates will be made at https://github.com/qnano/smfish.

### Statistical analyses

Data were visualized and analysed statistically using GraphPad Prism Version 8.3.1 (332). The distribution of a dataset was assessed with a Shapiro-Wilk test. All smFISH abundance data followed a Gaussian distribution. smFISH abundance was compared between two groups using an unpaired t-test. smFISH abundance between multiple groups was compared using a one-way ANOVA followed by Tukey’s post hoc test. Proportions of transcriptionally active soma were compared to transcriptionally inactive soma using a Chi-Square test. YFP positive and negative smFISH intensity distributions were compared with a two-sided Wilcoxon rank-sum test. YFP fluorescence intensity across treatments was compared using a one-way ANOVA for Gaussian distributed data, and a Kruskal-Wallis test for non-Gaussian distributed data. Statistical significance is defined as p<0.05.

