## Supplementary figures and images for "Dendritic localization of mRNA in *Drosophila* Mushroom Body Output Neurons"

### Supplemental Figure 1

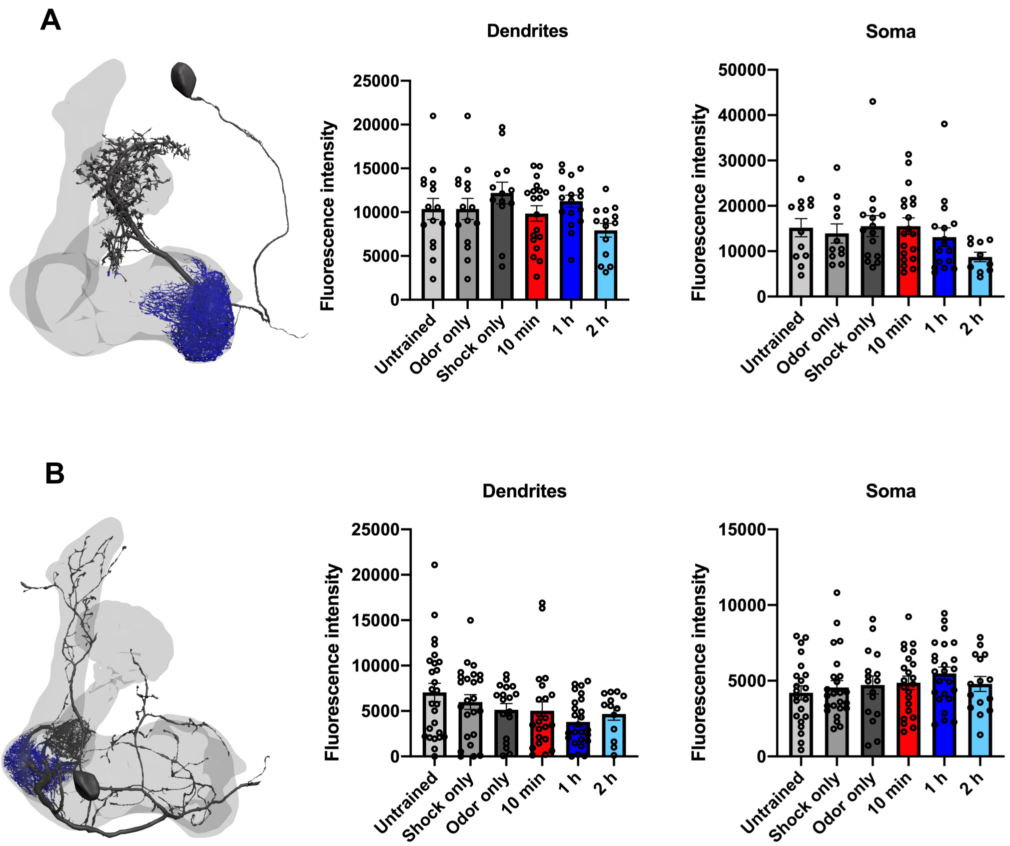
