## Supplementary material for "Dendritic localization of mRNA in *Drosophila* Mushroom Body Output Neurons": Key Resources Table

| REAGENT or RESOURCE | SOURCE | IDENTIFIER |
|---|---|---|
| Chemicals, Peptides and Recombinant Proteins | | |
| 20% v/v paraformaldehyde | ThermoFisher Scientific | Cat#15713S |
| RNase free 1x PBS | ThermoFisher Scientific | Cat#AM9625 |
| Triton X-100 | Sigma-Aldrich | Cat#T8787 |
| 20x RNase free SSC | ThermoFisher Scientific | Cat#AM9763 |
| Deionized formamide | ThermoFisher Scientific | Cat#AM9342 |
| 50% dextran sulphate | Millipore | Cat#S4030 |
| Vybrant DyeCycle Violet Stain | ThermoFisher Scientific | Cat#V35003 |
| Vectashield anti-fade mounting medium | Vector Laboratories | Cat#H-1000-10 |
| JF549-SNAPTag | (Grimm et al., 2015) | / |
| Mineral oil | Sigma-Aldrich | Cat#M5904 |
| 4-methocyclohexanol (98%) | Sigma-Aldrich | Cat#218405 |
| 3-octanol (99%) | Sigma-Aldrich | Cat#153095 |
| Experimental Models: Organisms/Strains | | |
| *D. melanogaster*: R66C08-Gal4 | Bloomington Drosophila Stock Center; (Owald et al., 2015) | RRID:BDSC_49412 |
| *D. melanogaster*: MB112c-Gal4 | Bloomington Drosophila Stock Center; (Perisse et al., 2016) | RRID:BDSC_68263 |
| *D. melanogaster*: UAS-myr::SNAPf | Bloomington Drosophila Stock Center | RRID:BDSC_58376 |
| *D. melanogaster*: CaMKII::YFP | Kyoto Stock Centre; (Lowe et al., 2014) | RRID:DGGR_115127 |
| *D. melanogaster*: PKA-R2::YFP | Kyoto Stock Centre; (Lowe et al., 2014) | RRID:DGGR_115174 |
| *D. melanogaster*: Ten-m::YFP | Kyoto Stock Centre; (Lowe et al., 2014) | RRID:DGGR_115131 |
| Software and Algorithms | | |
| FIJI | NIH; (Schindelin et al., 2012) | http://fiji.sc/ |
| MatLab R2019b | The Mathworks, Natick, MA | https://www.mathworks.com/products/matlab.html |
| GraphPad Prism 8 | GraphPad Software, La Jolla, CA | https://www.graphpad.com/scientific-software/prism/ |
| *Drosophila* brain smFISH analysis | Here | https://github.com/qnano/smfish |
| Blender | Blender Foundation, Amsterdam | https://www.blender.org |

| | | |
|---|---|---|
| NAVis 0.2.0 | (Bates et al., 2020) | https://pypi.org/project/navis/ |
