## Supplementary Methods for "Dendritic localization of mRNA in *Drosophila* Mushroom Body Output Neurons"

| Gene | Probe number | 5'-3' nucleotide sequence | Dye |
|---|---|---|---|
| *CaMKII* | 1 | GAAAAACGCGTACAGGCTGCTG | ATTO-633 |
| *CaMKII* | 2 | CTCTTCTTTGATGTCGTAATTG | ATTO-633 |
| *CaMKII* | 3 | ACAATTGAGAATGCTCCTTTTC | ATTO-633 |
| *CaMKII* | 4 | GTTGACTTTTGGACACATCTTT | ATTO-633 |
| *CaMKII* | 5 | ATTTTTGCAGCAAATTCAAAGC | ATTO-633 |
| *CaMKII* | 6 | TCTGGCAGTTAATTTTTTTGTA | ATTO-633 |
| *CaMKII* | 7 | TTCACGTTCCAGTTTTTGAAAG | ATTO-633 |
| *CaMKII* | 8 | TATGTTGGGATGGTGTAACTTT | ATTO-633 |
| *CaMKII* | 9 | AACAAGGTAGTGATAGTTCTCC | ATTO-633 |
| *CaMKII* | 10 | CTTCTGAATAAAATTCGCGTGC | ATTO-633 |
| *CaMKII* | 11 | GTTGAATACAATGTGATGCATC | ATTO-633 |
| *CaMKII* | 12 | AGTGATTGACCGATTCCAATAT | ATTO-633 |
| *CaMKII* | 13 | AATTCTCTGGTTTCAGATCTCG | ATTO-633 |
| *CaMKII* | 14 | CCTTAGCCTTACTAGCTAATAG | ATTO-633 |
| *CaMKII* | 15 | TAGACCAAAGTCAGCGAGTTTC | ATTO-633 |
| *CaMKII* | 16 | ATGATCGCCTTGAACTTCAATG | ATTO-633 |
| *CaMKII* | 17 | AATACCTCAGGAGACAGATATC | ATTO-633 |
| *CaMKII* | 18 | GTAAAGAATAACTCCACATGCC | ATTO-633 |
| *CaMKII* | 19 | AAGGGTGGATAACCGACAAGAA | ATTO-633 |
| *CaMKII* | 20 | CTTTTATCTGCGAGTACAGTCG | ATTO-633 |
| *CaMKII* | 21 | CATTCTGGAGACGGATAATCAT | ATTO-633 |
| *CaMKII* | 22 | GATTTTTAGCTTCTGGAGTAAC | ATTO-633 |
| *CaMKII* | 23 | GATTAACGGTTAGCATTTGATT | ATTO-633 |
| *CaMKII* | 24 | CAGCTGCAGTTATTCGTTTATT | ATTO-633 |
| *CaMKII* | 25 | TGACAAATCCACGGATGTTTTA | ATTO-633 |
| *CaMKII* | 26 | GCGCATTAAATTTCTTGAGACA | ATTO-633 |
| *CaMKII* | 27 | AACATTGTCGTAAGTATGGCTC | ATTO-633 |
| *CaMKII* | 28 | TTCTGCTCGAAAAATTTCTGGT | ATTO-633 |
| *CaMKII* | 29 | CTTCACCTTTTTTAGTTATCAT | ATTO-633 |
| *CaMKII* | 30 | GATGAATCGGTTGATTCTTTGA | ATTO-633 |
| *CaMKII* | 31 | TTTTATGTCATCGTCTTCAAGA | ATTO-633 |
| *CaMKII* | 32 | TATTATTTCTTGTCGTCTAGCT | ATTO-633 |
| *CaMKII* | 33 | TTCAATCAACTGCTCAGTTATC | ATTO-633 |
| *CaMKII* | 34 | ATCAAAGTCGCCACTGTTAATT | ATTO-633 |
| *CaMKII* | 35 | TTAGATGCGGATCACATATTTT | ATTO-633 |
| *CaMKII* | 36 | ATCAATTCCTTCTACAAGGTTA | ATTO-633 |
| *CaMKII* | 37 | GTTTTTACCAAGAACATTTTCA | ATTO-633 |
| *CaMKII* | 38 | AATGGTTGTGTTTATAGCTTTG | ATTO-633 |
| *CaMKII* | 39 | CAAGTAAGTGCACATGAGGATT | ATTO-633 |
| *CaMKII* | 40 | CGTAAGTCTCACATAGGCAATG | ATTO-633 |
| *CaMKII* | 41 | ATGTCCTTGCTTATCAATGTAC | ATTO-633 |
| *CaMKII* | 42 | AACATTCTGCCATTTGTTATCG | ATTO-633 |
| *CaMKII* | 43 | GGCAGATGCACTTCGATGAAAA | ATTO-633 |
| *CaMKII* | 44 | AAAATCGAATGTAGTTGCTCCG | ATTO-633 |

| Gene | Probe number | 5'-3' nucleotide sequence | Dye |
|---|---|---|---|
| *nAChRα1* | 1 | ATACAGCTGCGAATAGCACG | ATTO-633 |
| *nAChRα1* | 2 | GGTGGCAAAGTGTAATGCTA | ATTO-633 |
| *nAChRα1* | 3 | GGATGAGGCGATTGTAGTTG | ATTO-633 |
| *nAChRα1* | 4 | ATCTTGACGGTGAGACGGTC | ATTO-633 |
| *nAChRα1* | 5 | GATTCACATCGATCAGCTGG | ATTO-633 |
| *nAChRα1* | 6 | CCAGACATTGGTTGTCATAA | ATTO-633 |
| *nAChRα1* | 7 | ATTTATAGTCGTTCCATTCC | ATTO-633 |
| *nAChRα1* | 8 | ATAGTCATCCGGATTCCATT | ATTO-633 |
| *nAChRα1* | 9 | GAGGGAACGTGCAAAGTGTC | ATTO-633 |
| *nAChRα1* | 10 | ATATCCGGTAGCCATATATG | ATTO-633 |
| *nAChRα1* | 11 | ATCGGCGTTGTTATAGAGCA | ATTO-633 |
| *nAChRα1* | 12 | GCTTTTGTCATTATTGTCAC | ATTO-633 |
| *nAChRα1* | 13 | TTTGCCCGTGTGATGAAGAA | ATTO-633 |
| *nAChRα1* | 14 | TCTGCTCGTCGAAGGGAAAG | ATTO-633 |
| *nAChRα1* | 15 | CAGGATCCGAACTTCATGAA | ATTO-633 |
| *nAChRα1* | 16 | CACCATGTAACCATCGTAGG | ATTO-633 |
| *nAChRα1* | 17 | TCTGCTTCAAGTGCCTCAAG | ATTO-633 |
| *nAChRα1* | 18 | ACTCCACGGAGATGTAGTAG | ATTO-633 |
| *nAChRα1* | 19 | AGCTGTAGAACTTCTCGTTC | ATTO-633 |
| *nAChRα1* | 20 | GGTTGAACACTATGTCCAGA | ATTO-633 |
| *nAChRα1* | 21 | TGTAGAAGAGCGTCTTCCGG | ATTO-633 |
| *nAChRα1* | 22 | CAGGGTATGATCAGGTTGAC | ATTO-633 |
| *nAChRα1* | 23 | CCAGAACGGACAGGAACGAG | ATTO-633 |
| *nAChRα1* | 24 | TGATGCAGAGCGAGATCTTC | ATTO-633 |
| *nAChRα1* | 25 | CAGCAGGAGGAAGAACACGG | ATTO-633 |
| *nAChRα1* | 26 | TGAAGAGCAGATACTTTCCC | ATTO-633 |
| *nAChRα1* | 27 | ACGACAGAGAGCGTGACCAG | ATTO-633 |
| *nAChRα1* | 28 | TCACATTGAGCACGGCAATG | ATTO-633 |
| *nAChRα1* | 29 | TGCGTGACAGGGGATCTAAA | ATTO-633 |
| *nAChRα1* | 30 | GCAGGATCTGGATGAAGAGG | ATTO-633 |
| *nAChRα1* | 31 | CAGGTGATAGACATCGGTGA | ATTO-633 |
| *nAChRα1* | 32 | TGACAAACTTGTCCACATCC | ATTO-633 |
| *nAChRα1* | 33 | CGCTGAAACGCTTCGAATCG | ATTO-633 |
| *nAChRα1* | 34 | AAGCGGGTAGTGCTGGAATG | ATTO-633 |
| *nAChRα1* | 35 | GCAGCCAAATCGAAGCGATG | ATTO-633 |
| *nAChRα1* | 36 | CGAAACAGTGGGCGCTAATC | ATTO-633 |
| *nAChRα1* | 37 | GACGGGCTGAACAGATCGTC | ATTO-633 |
| *nAChRα1* | 38 | GACTGATGTCTCCGTTGAGG | ATTO-633 |
| *nAChRα1* | 39 | CTTCTCGAACGTAGGACTGA | ATTO-633 |
| *nAChRα1* | 40 | TTCGATGGTCTTCTCCATTT | ATTO-633 |
| *nAChRα1* | 41 | CCACACTCTCAAACTTATCC | ATTO-633 |
| *nAChRα1* | 42 | ATACCATGGCAACGTACTTC | ATTO-633 |
| *nAChRα1* | 43 | GATCGCGAAGATCCACAGAA | ATTO-633 |
| *nAChRα1* | 44 | CTGCAGGATAATCAGCGCTG | ATTO-633 |
| *nAChRα1* | 45 | TATATCGATCGGCTGCGATT | ATTO-633 |
| *nAChRα1* | 46 | ATCTTGAGCAGCTCGAACTT | ATTO-633 |

| Gene | Probe number | 5'-3' nucleotide sequence | Dye |
|---|---|---|---|
| *YFP* | 1 | CGGTGAACAGCTCCTCGC | Quasar-670 |
| *YFP* | 2 | GACCAGGATGGGCACCAC | Quasar-670 |
| *YFP* | 3 | GTTTACGTCGCCGTCCAG | Quasar-670 |
| *YFP* | 4 | CCGGACACGCTGAACTTG | Quasar-670 |
| *YFP* | 5 | TTGCCGTAGGTGGCATCG | Quasar-670 |
| *YFP* | 6 | CCGGTGGTGCAGATGAAC | Quasar-670 |
| *YFP* | 7 | AGGGTGGTCACGAGGGTG | Quasar-670 |
| *YFP* | 8 | AAGCACTGCACGCCGTAG | Quasar-670 |
| *YFP* | 9 | ATGTGGTCGGGGTAGCGG | Quasar-670 |
| *YFP* | 10 | TGAAGAAGTCGTGCTGCT | Quasar-670 |
| *YFP* | 11 | ACGTAGCCTTCGGGCATG | Quasar-670 |
| *YFP* | 12 | AGAAGATGGTGCGCTCCT | Quasar-670 |
| *YFP* | 13 | TCTTGTAGTTGCCGTCGT | Quasar-670 |
| *YFP* | 14 | TCGAACTTCACCTCGGCG | Quasar-670 |
| *YFP* | 15 | TCGATGCGGTTCACCAGG | Quasar-670 |
| *YFP* | 16 | TGAAGTCGATGCCCTTCA | Quasar-670 |
| *YFP* | 17 | CAGGATGTTGCCGTCCTC | Quasar-670 |
| *YFP* | 18 | TTGTACTCCAGCTTGTGC | Quasar-670 |
| *YFP* | 19 | AGACGTTGTGGCTGTTGT | Quasar-670 |
| *YFP* | 20 | CTGCTTGTCGGCCATGAT | Quasar-670 |
| *YFP* | 21 | TTCACCTTGATGCCGTTC | Quasar-670 |
| *YFP* | 22 | CGATGTTGTGGCGGATCT | Quasar-670 |
| *YFP* | 23 | TAGTGGTCGGCGAGCTGC | Quasar-670 |
| *YFP* | 24 | CGATGGGGGTGTTCTGCT | Quasar-670 |
| *YFP* | 25 | TTGTCGGGCAGCAGCACG | Quasar-670 |
| *YFP* | 26 | GACTGGGTGCTCAGGTAG | Quasar-670 |
| *YFP* | 27 | GTTGGGGTCTTTGCTCAG | Quasar-670 |
| *YFP* | 28 | ACCATGTGATCGCGCTTC | Quasar-670 |
| *YFP* | 29 | CGGTCACGAACTCCAGCA | Quasar-670 |
| *YFP* | 30 | TCCATGCCGAGAGTGATC | Quasar-670 |

| Gene | Probe number | 5'-3' nucleotide sequence | Dye |
|---|---|---|---|
| *nAChRα5* | 1 | CAACTTCAGTCAGTTTCAGTTG | ATTO-633 |
| *nAChRα5* | 2 | ACAGTCTTATGTTTGTTGGTTG | ATTO-633 |
| *nAChRα5* | 3 | TTCTTGATATCTGTTTCGTTTG | ATTO-633 |
| *nAChRα5* | 4 | TTTTGAAGGGAGGCATGCTAAG | ATTO-633 |
| *nAChRα5* | 5 | AATACTGAACTCATTGTCGCTG | ATTO-633 |
| *nAChRα5* | 6 | TACTCTATCATGTCTCGATATC | ATTO-633 |
| *nAChRα5* | 7 | GCAAAGCTGCCTAAATAGGAAA | ATTO-633 |
| *nAChRα5* | 8 | GAGTGTTTATTAAGTCCGTTTA | ATTO-633 |
| *nAChRα5* | 9 | CTAGGCAAACTTTAGCAGATAA | ATTO-633 |
| *nAChRα5* | 10 | AACAGTCTCTTTTCATGATATC | ATTO-633 |
| *nAChRα5* | 11 | TTATAGGGATCCAAAAGATCGT | ATTO-633 |
| *nAChRα5* | 12 | ATTCATTGAGAACGGGACGTTC | ATTO-633 |
| *nAChRα5* | 13 | ACCAAAGCTTAATTGTAACGGG | ATTO-633 |
| *nAChRα5* | 14 | CCACATCGATAATTTGCATTAA | ATTO-633 |
| *nAChRα5* | 15 | TGACTAGCAATTGATTTTTCTC | ATTO-633 |
| *nAChRα5* | 16 | ACTCCAGTTTTAACCACACATT | ATTO-633 |
| *nAChRα5* | 17 | AACGTATAGACACGAGCCGTTG | ATTO-633 |
| *nAChRα5* | 18 | CTTGCACGTCGACTTGAAGATC | ATTO-633 |
| *nAChRα5* | 19 | TCTTGTAATTGTAAATCCAGCT | ATTO-633 |
| *nAChRα5* | 20 | TTGAGCACGTAACTGCTGATAT | ATTO-633 |
| *nAChRα5* | 21 | AGCAGTTGTAATAGATTTCGTT | ATTO-633 |
| *nAChRα5* | 22 | AAGGTGATGTCTATATAGGGTT | ATTO-633 |
| *nAChRα5* | 23 | AGAAATAGTACAGTGTTCGTCG | ATTO-633 |
| *nAChRα5* | 24 | GCAATCAGTACACAAGGTATGA | ATTO-633 |
| *nAChRα5* | 25 | AGTGATAATTTTTCACCCGAAT | ATTO-633 |
| *nAChRα5* | 26 | AGCGAGAGCAAGATGGTAACAC | ATTO-633 |
| *nAChRα5* | 27 | GGCAACCATATTCAGAAACACG | ATTO-633 |
| *nAChRα5* | 28 | TGAAATATGTACCCAGCAATGG | ATTO-633 |
| *nAChRα5* | 29 | AAGCTACCATAAACATTATGCA | ATTO-633 |
| *nAChRα5* | 30 | TTAAAATCGTTGACACAACGGA | ATTO-633 |
| *nAChRα5* | 31 | GTATCAGCATTTCGATGATGAT | ATTO-633 |
| *nAChRα5* | 32 | ACAAAAACACGATGCGTATCCA | ATTO-633 |
| *nAChRα5* | 33 | CGACTCATTCGCAATATCCATG | ATTO-633 |
| *nAChRα5* | 34 | AAGTCATCATCGATGTCTAGTA | ATTO-633 |
| *nAChRα5* | 35 | CCATAAACCGTGCGATAGAAAG | ATTO-633 |
| *nAChRα5* | 36 | TTTGATGCACGTATGATGGGTG | ATTO-633 |
| *nAChRα5* | 37 | ACCTAATTCATATTCAGTTGAT | ATTO-633 |
| *nAChRα5* | 38 | TCAGTTATAAAGCGAATTTCCT | ATTO-633 |
| *nAChRα5* | 39 | TCATTGCACTCGTCATCTTTAC | ATTO-633 |
| *nAChRα5* | 40 | AATATGATAAGGCACAGTCTGT | ATTO-633 |
| *nAChRα5* | 41 | ACAGCTATTGTGGCTAATATTG | ATTO-633 |
| *nAChRα5* | 42 | CGAGACAATAATATGTGGTGCT | ATTO-633 |

| Gene | Probe number | 5'-3' nucleotide sequence | Dye |
|---|---|---|---|
| *nAChRα6* | 1 | AACAGGACAAACAGCGACAGC | ATTO-633 |
| *nAChRα6* | 2 | TTTCTTTAATTATCGCCAGAA | ATTO-633 |
| *nAChRα6* | 3 | CTTTTCATGAGGTCCTTGACA | ATTO-633 |
| *nAChRα6* | 4 | TATTGTAGGTGGACAGCAGAT | ATTO-633 |
| *nAChRα6* | 5 | CAGCGTCAGTCCGAACTTAAC | ATTO-633 |
| *nAChRα6* | 6 | TTCATCCACGTCGATGATCTG | ATTO-633 |
| *nAChRα6* | 7 | TTTGTGGTCAGAATCTGATTC | ATTO-633 |
| *nAChRα6* | 8 | TCTCGTCCAAATTTAACCACG | ATTO-633 |
| *nAChRα6* | 9 | ATTCGTTATGAGAAGCTGATT | ATTO-633 |
| *nAChRα6* | 10 | TTCCACTCCAACGAAAGCCAA | ATTO-633 |
| *nAChRα6* | 11 | TCATTCCAGCGCAGATTGTAG | ATTO-633 |
| *nAChRα6* | 12 | CGTGATTCGTAGATCCTTGAC | ATTO-633 |
| *nAChRα6* | 13 | CTGTTGTACATGAGCACGTCG | ATTO-633 |
| *nAChRα6* | 14 | CACAATGTTGGTGTGATACGT | ATTO-633 |
| *nAChRα6* | 15 | AGACAACTGCCGTTATGTTTG | ATTO-633 |
| *nAChRα6* | 16 | ATGTGCTCTTGAAGATACCAG | ATTO-633 |
| *nAChRα6* | 17 | GAACCACGTGATGTCTATCTT | ATTO-633 |
| *nAChRα6* | 18 | TCGCAATGTTGGTCATCAAAT | ATTO-633 |
| *nAChRα6* | 19 | AAGTCCAACTACCGAATTTCA | ATTO-633 |
| *nAChRα6* | 20 | CAAATCCAACTGATTTCCATC | ATTO-633 |
| *nAChRα6* | 21 | CCTCCATCTTCGGAATTCAAA | ATTO-633 |
| *nAChRα6* | 22 | CGCCATTTGTTATGAAATCGG | ATTO-633 |
| *nAChRα6* | 23 | CGTAGACTATCGTATTCTTCT | ATTO-633 |
| *nAChRα6* | 24 | GGTGATATCGACATATGGTTC | ATTO-633 |
| *nAChRα6* | 25 | ACGGCGACGAATTTGTATAGT | ATTO-633 |
| *nAChRα6* | 26 | ATTAGCACACATGGCACAATT | ATTO-633 |
| *nAChRα6* | 27 | GAATCCGGCGGCAATGTGAAG | ATTO-633 |
| *nAChRα6* | 28 | AATTGTAACTCCCAGCGTCAG | ATTO-633 |
| *nAChRα6* | 29 | AGAAACACTGTGAGCGATAGA | ATTO-633 |
| *nAChRα6* | 30 | GCAATGTCTCAGCTACAAGGT | ATTO-633 |
| *nAChRα6* | 31 | AGGGGATTGCATCAGATACTT | ATTO-633 |
| *nAChRα6* | 32 | GCAATTGAAGTAGGTGCCTAA | ATTO-633 |
| *nAChRα6* | 33 | GACGAGGCGACCATGAACATG | ATTO-633 |
| *nAChRα6* | 34 | TAGTTGAGCACCACTACTGTC | ATTO-633 |
| *nAChRα6* | 35 | AGCCATTGTAGGAAAACGGAC | ATTO-633 |
| *nAChRα6* | 36 | TTGTTTTGCGTGTAATCTTGC | ATTO-633 |
| *nAChRα6* | 37 | CCTTCATGCGATTGCTTAATA | ATTO-633 |
| *nAChRα6* | 38 | TCGTCGATGTCGAGGACATTG | ATTO-633 |
| *nAChRα6* | 39 | CAGATATTGTGTGCCGGAAGT | ATTO-633 |
| *nAChRα6* | 40 | GAAGATCTTTGTGATTGCAGC | ATTO-633 |
| *nAChRα6* | 41 | CGCGCCGTAATAAATTGCAAT | ATTO-633 |
| *nAChRα6* | 42 | AACTTCCAATCGCCGATCAAT | ATTO-633 |
| *nAChRα6* | 43 | AAATCTATCCACAACCATTGC | ATTO-633 |
| *nAChRα6* | 44 | CCGTTGCAATAATCGTGAAGA | ATTO-633 |
| *nAChRα6* | 45 | TATTGCACGATTATGTGCGGA | ATTO-633 |
